# Measuring Orangutan nest structure using Unmanned Aerial Vehicle (UAV) and ImageJ

**DOI:** 10.1101/365338

**Authors:** Salniza Akmar Kamaruszaman, Nik Fadzly, Aini Hasanah Abd Mutalib, Aidy M. Muslim, Sri Suci Utami Atmoko, Mashhor Mansor, Asyaf Mansor, Nadine Rupert, Rahmad Zakaria, Zarul Hazrin Hashim, Amir Shah Ruddin Md Sah, Fadhirul Fitri Jamsari, Nur Munira Azman

## Abstract

The nest is one of the crucial elements in orangutan daily activities. Previously, most of the nest structure studies were done manually by estimating measurement directly from visual observation. However, using the latest unmanned aerial vehicle (UAV) technology, we can reduce the workforce, time and energy while simultaneously ensuring the safety of the researcher conducting nest structure analysis. We recorded 49 pictures of orangutan nests at Sepilok Orangutan Rehabilitation Centre (SORC) using UAV (DJI Phantom 3 Quadcopter). The nest structure (length, depth, and width) was digitally measured by using ImageJ. Most of the nests were built at a strong, stable, and comfortable position at the top of the tree. Most orangutans chose *Eusideroxylon zwageri* to build nest compared to other tree species because of the strong and durable wood characteristic which would create a sturdy, strong and comfortable nest. We propose the use of drone with digital image analysis could provide a more accurate, less time consuming and safe method for studying orangutan nest structure.

## Introduction

Arboreal great apes especially orangutans need to master the nest building skill together with other skills such as climbing, foraging and being able to identify their natural predators [1–4]. The nest building is a skill inherited through observation on the mother’s or other adults’ nesting practices [2, 3, 5, 6]. Nests served as a "bed" for resting and sleeping, hiding from danger or predator, as well as for a better thermoregulation [4, 5, 7].

Previously, most of the orangutans’ nest studies were based on ecological aspects such as density estimation, distribution and population of orangutans in a habitat, nesting preferences and mechanisms and material to build a nest and its decaying rates [7–12]. Nest measurement and materials used are evaluating characteristics of the nest building skill of an individual and the quality of the nest built [6, 13]. However, researchers must conduct rigorous, time and energy consuming techniques such as tree climbing to study the orangutan’s nest structures. Even though direct nests measurement by climbing techniques might provide a more accurate reading, a new approach using Unmanned Aerial Vehicle (UAV) has been developed for nest survey and observation during this decade [7, 11, 13–15].

The application of UAV or drones technology was once limited to the military. However, the usage of drones has spread widely and no longer exclusive to the military. Drones are used in civilian use such as monitoring, transportation for goods delivery as well as site inspections [16, 17]. These drones have the capability to capture images through rapid data acquisition, which is an advantage for scientific research especially for animals conservations [8, 11, 16–19]. This study aims to determine the variation of nest structure quantitatively via image analysis through utilization of drones.

## Materials and Method

We obtained the permission to conduct the study from Sabah Biodiversity Centre (SaBC) with the support of Sabah Wildlife Department (SWD) and Sabah Forestry Department (SFD). The study was done at 5°51′51.82″N, 107°56′55.72″E; Sepilok Orangutan Rehabilitation Center (SORC), Sandakan Sabah within 6 months from January 2016 to April 2017. SORC is located within Kabili-Sepilok Forest Reserved with an area of 4294 hectares with more than 200 Orangutans (Fig 1).

**Fig 1.**
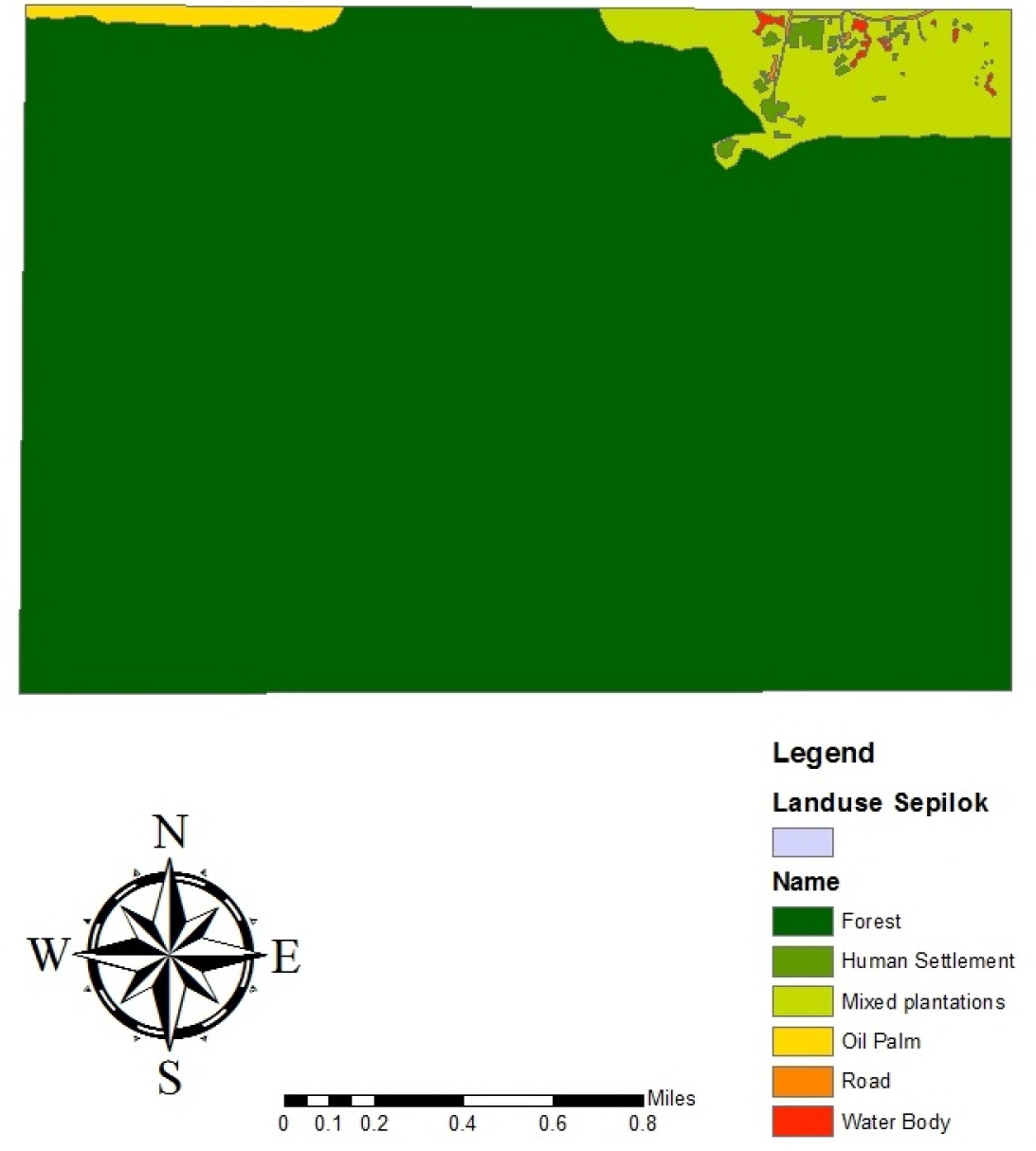
The area of study; Sepilok Orangutan Rehabilitation Centre (Map by Aini Hasanah Abd Mutalib).

We tested a new experimental design by adapting to the nest description outlined by Samson and Hunt (6). We utilized the use of Unmanned Aerial Vehicle (UAV) to capture the orangutan nests images [11, 20]. We used the DJI Phantom 3 Professional fitted with a 12-megapixel camera (f/2.8 lens 94^0^ field view). Image capture process was done without the presence of the orangutans during daytime for safety purposes (for the researchers and the orangutan). Our skilled licensed drone pilots managed to fly the drone as close as within 1 to 3 meters of the nest. The side and top view of the nest were recorded at a screen resolution of 72 dpi (dot per inch) which equals to 300 dpi in print resolution. The digital scale measurement of the nests’ structure was estimated by using leaf samples of the tree where the nests were recorded. We used a slingshot to obtain the leaf samples from the nearest branch of the nest. The leaf samples were measured by using a ruler to obtain the length. Ten samples of the leaves were measured and we used the average value. In the ImageJ software, we used the measured length as the digital scale for the overall image analysis. ImageJ software was used to measure the nest length, width, and depth. The digital measurement of the nest was illustrated in Fig 2.

**Fig 2.**
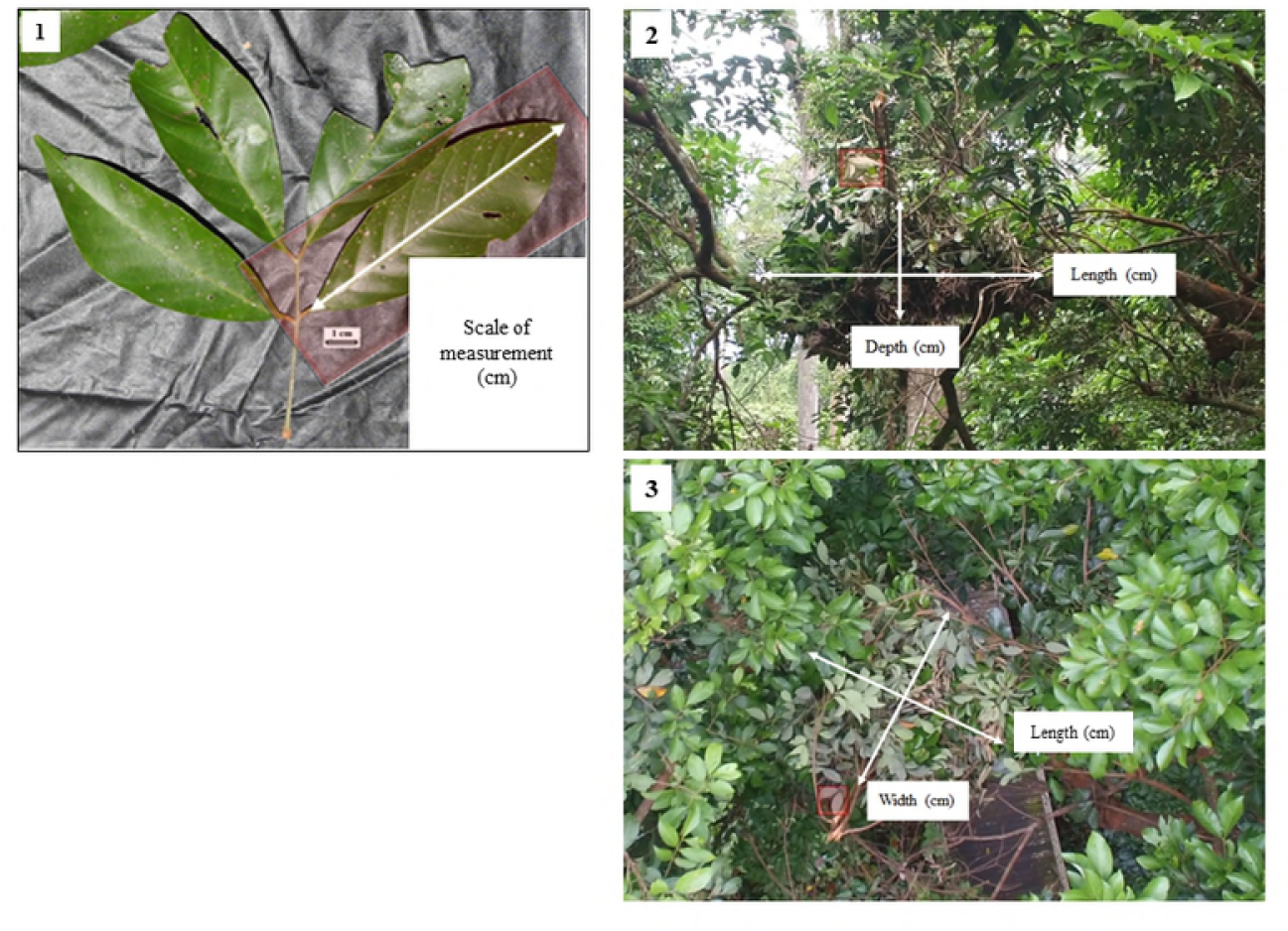
The mechanism of nest measurement using the ImageJ analysis was modified from van Casteren *et al*.,[7], **1.** Leaf sample (red boxes) was measured and set as digital scale (cm). **2.** The length of the nest was measured, the depth of the nest was taken from the centre of the nest. **3.** The nest width was measured perpendicular to the length.

Other ecological parameters (nest class and position, canopy cover, tree height, nest height, and tree species) were also recorded through direct observation. Nest class and position were classified based on Utami Atmoko and Arif Rifqi (4) and Prasetyo, Utami (1). The nest canopy cover was classified directly through observation in the field. The nest was categorized as closed nest if there was the presence of tree canopy, vegetation, or any obstacle above the nest. A nest was considered as an open nest with the absence of any obstacles that prevent direct sunlight to the nest. The tree and nest height was either measured by using a clinometer (Suunto PM-5, PM-5/1520) or estimated through observation. The tree species were identified by the available tree id tags and later reconfirmed by the SFD’s staff. The data were analyzed by using statistical software JMP 10. The variables that were not normally distributed were transformed using log to meet the condition of normal distribution for parametric analyses. If normality could not be achieved, we proceeded to use non-parametric analyses.

## Results

We recorded a total of 49 nests. Fig 3a shows most of the nest were built at tree branch or position 2 with 25 nests count followed by position 3 (16 nests) and position 1 (8 nests). Fig 3b shows that 27 of the nests recorded were from class 3 followed by class 1 (11 nests), class 2 (10 nests) and class 4 (1 nest). In this study, we do not encounter any nest with position 4.

**Fig 3.**
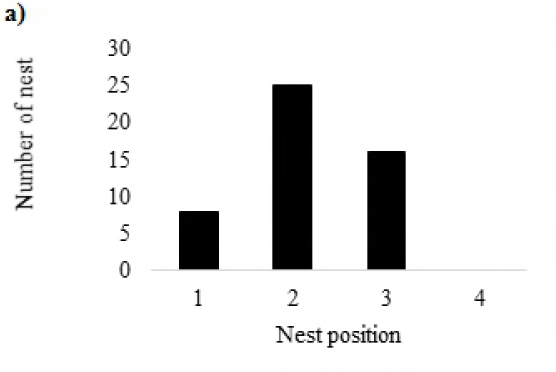
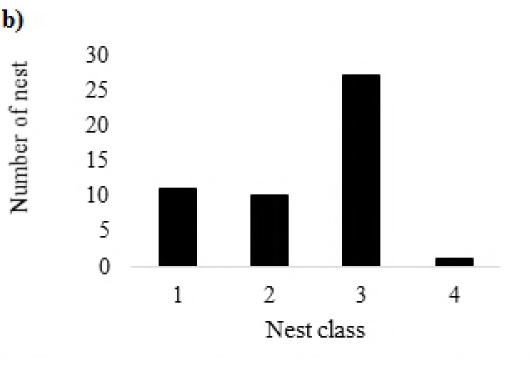
The bar graph of number of nests built by orangutans based on **a)** nest position and **b)** nest class.

The average nest length, width and depth recorded were 87.323 ± 29.472 cm, 59.889 ± 18.313 cm and 36.666 ± 16.009 cm, respectively. The average height of the tree, and the height of nest were 16.166 ± 7.686 m, and 12.176 ± 6.866 m, respectively. 65.63% of the nests were open nest and 34.63% of the nests were closed nest (Fig 4).

**Fig 4.**
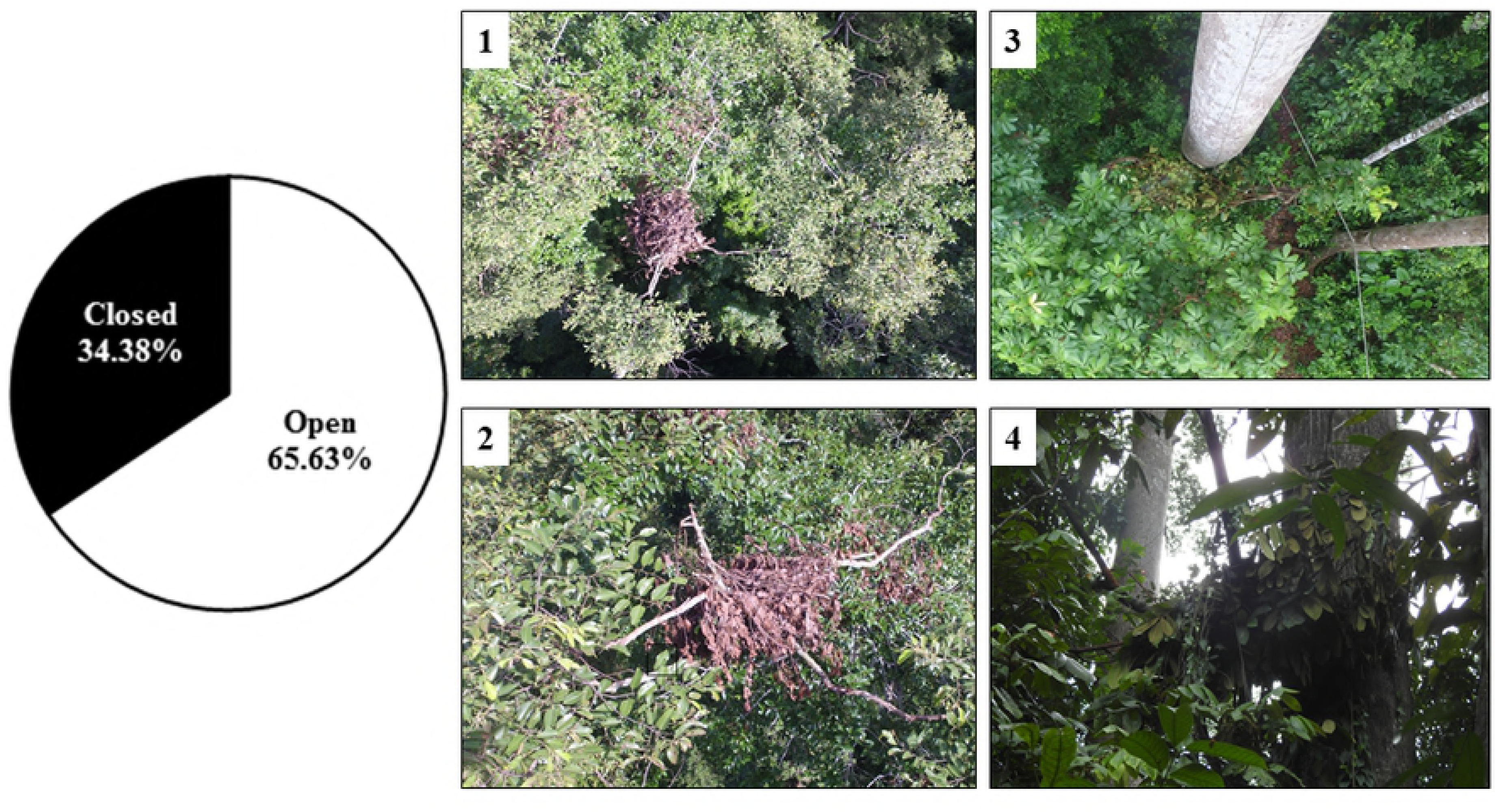
The pie chart shows the percentage nests built according to nest cover. Inset picture 1 and 2 show an open nest, inset picture 3 and 4 show a closed nest, from the top and side view.

There was no significant difference for nest position according to nest length (Kruskall-Wallis, χ^2^ = 2.138, df = 2, P = 0.343) and depth (Kruskall-Wallis, χ^2^ = 2.108, df = 2, P = 0.349). There was also no significant difference for the nest width based on the nest position (ANOVA, F (2,77) = 1.839, P = 0.166). There was no significant difference for the nest class based on the nest length (Kruskall-Wallis, χ^2^ = 4.304, df = 3, P = 0.231).

There is a significant difference of nest depth based on the nest class (Kruskall-Wallis, χ^*2*^ = 13.408, df = 2, P = 0.001) (Fig 5). The depth of nest decreased with the nest class. Class 3 nest recorded the thinnest depth (34.560 ± 3.613 cm) compared to nest from class 1 (52.200 ± 2.032 cm and class 2(57.000 ± 2.430 cm). There was no significant difference for nest class according to the nest width (F (3,77) = 2.187, P = 0.097. Tree height and nest height showed a strong positive and significant correlation (r_s_ =0.7867, p=0.0001) (Fig 6).

**Fig 5:**
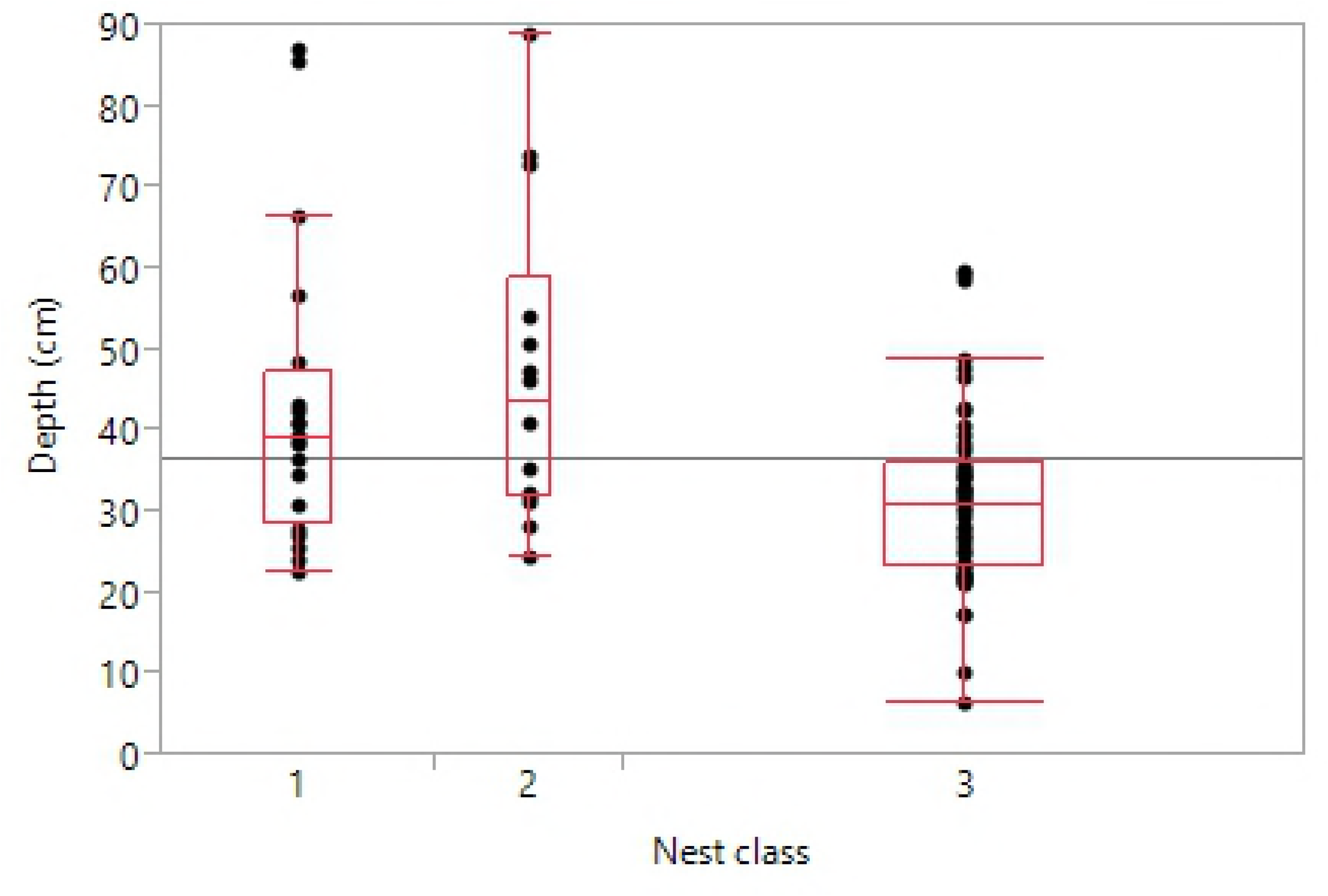
The nest depth based on the nest class distribution.

**Fig 6:**
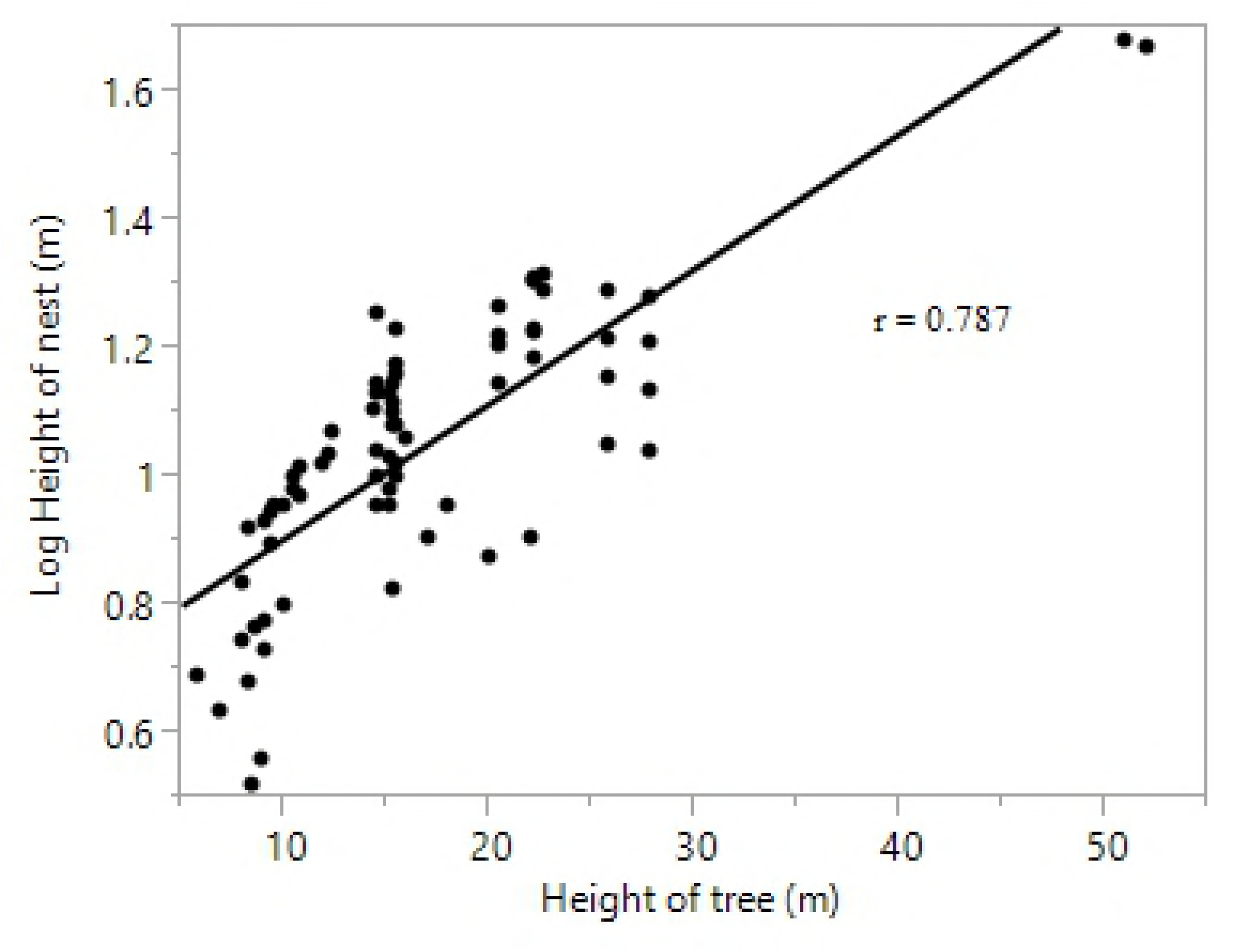
The correlation between a) height (log) of tree and height of nest

From the 49 nests recorded, 46.94% (23 nests) were found on *Eusideroxylon zwageri*, 20.41% (10 nests) on *Nephelium rambutan-ake*, 8.16% (4 nests) on *Litsea* sp. and *Syzygium rejangense*, 4.08% (2 nests) *Pometia pinnata* and *Pentace laxiflora* and 2.04% (1 nest) on *Koordersiodendron pinnatum, Knema latifolia, Shorea johorensis*, and *Nephelium lappaceum*. (Fig 7)

**Fig 7:**
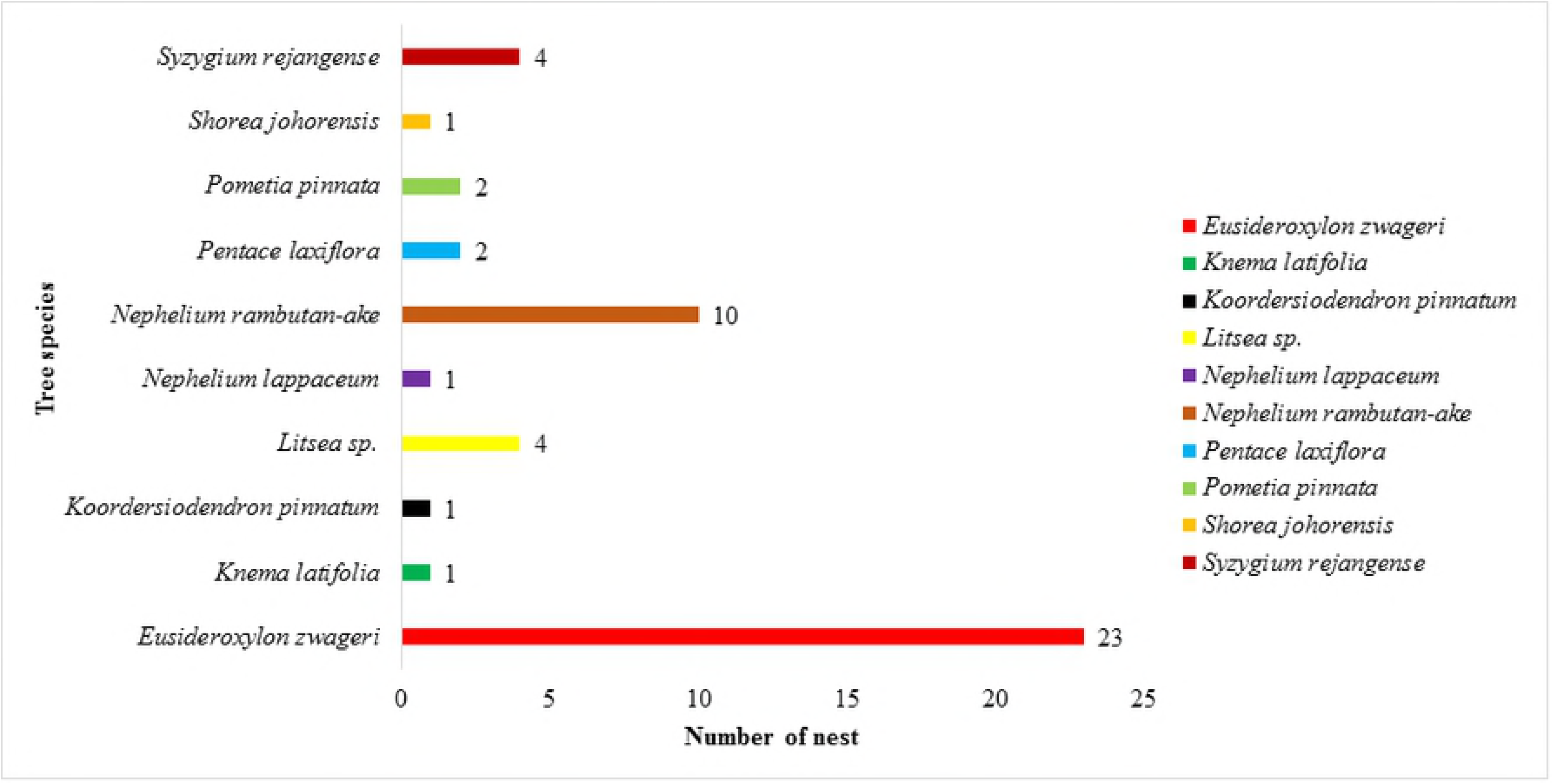
The frequency of tree species used for nesting by orangutan.

## Discussion

The aim of this study was to determine the variation of nest structure quantitatively using advance drone and image analysis software. We aim to promote our method as a new approach in nest measurement. Rayadin and Saitoh (13) recorded the nest average of length measurement of 114.5 cm. The average nest length for small juvenile to adult flange male was 64.1 cm to 139.3 cm. In this study, the average length of nest was 87.32 cm, which is within the range of the previous studies. However, we could not identify the individual that built the nest since we did not follow any orangutans. We were only allowed to use the drone when the orangutans had left the nest, for safety purposes. However, through observation we noted that juveniles and adults of both sex frequented the nests. We did not observe any adult flange male near our observed nests. Flanged male individual tend to avoid crowd areas, therefore their nest might be secluded.

Most of the nests at our study site were built on top of the tree with open nest cover. Ancrenaz, Calaque (12) reports that most of the orangutans build their nest on top of a single tree. Orangutans tend to choose higher trees to nest on regardless the age groups and sex [13, 21]. The trees in our study sites averages at 16.60 m where the top position might offer a stable and balanced position. Orangutans might have avoided small trees as they preferred strong and sturdy branches to support their concave shape nests. This is similar to chimpanzee nest selection preference, in which they preferred sturdy, strong and comfortable spot to build nest [1, 5–7].

Our results showed no significant differences between the nest sizes cumulatively. However, several studies have shown that nests size increases with the age group. This is related to the body size where adult flange male with large body size built larger nest compare to the juvenile with a small body [7, 13]. In our study site, since we did not detect any adult flanged male nesting and most of the nests that we sampled were frequented by orangutans within a similar age group and body sizes. There is a possibility that a larger sample size could provide more information in the future.

The significant correlation between tree height and nest height supports the hypothesis which orangutans were more likely to build the nest at the very top of the tree. The decaying process might also have contributed to the significant difference between nest class and nest depth [10]. Class 3 is considered as the second last stage of nest decay. Orangutans usually repairs certain nest spots that are their favorite, which is why nest class 1 and 2 is frequently used. However, the decay rate or factors affecting the decay process and decay process related to nest class quality were not recorded in this study. There is also the possibility that the depth of the nest was influenced by the orangutan body weight. Orangutan would position themselves in the middle of the nest and their body weight would depress the whole nest. Unfortunately, since we did not specifically track individual orangutans to a specific nest, we could not confirm this effect. There was also no assessment related to the animal physical measurement such as weight and body size as we were not allowed to physically touch the orangutans.

Orangutan most likely avoided building nest on fruit trees as precautions steps from other individuals especially flange males or other frugivorous animals. However, with limited availability of fruit, there were cases where an individual would select the non-fruiting tree to build their nest [5, 21, 22]. Orangutans do have individual preferences in where to rest and sleep [4, 5]. Orangutans shows intelligence in nest building by choosing the best materials from hard and durable tree species such as *E. zwageri* tree (commonly known as a Belian tree). Compared to other tree species, the Belian tree would provide the orangutans a sturdy and strong nest to support their weight. This valuable timber is widely used by human to produce furniture, medicinal purposes as well as in traditional rituals [23–26]. The frequent usage of this tree species also indicates that orangutans chose their nesting site based on strong and sturdy tree more for their comfort [27].

In natural settings, orangutans are solitary and have the tendency to avoid crowd and predators including human [4, 28, 29]. In this study, we noticed that most of the orangutans have become habituated with the human presence since most of the nests recorded were located near the feeding platforms and boardwalk; a place where the visitors could access and observe the orangutan’s activities. We also suspected that this phenomenon was also due to food availability since the food will be given to them every 10 a.m. and 3 p.m. regardless to the presence of visitors.

However, we must highlight the fact that the orangutans in Sepilok were in a rehabilitation program which indirectly means that they were already familiar with human presence since they have been exposed to human care. To release them into the wild, the juvenile orangutans were released in the rehabilitation area within the 4,294-hectare forest. This was the next step in preparing the orangutans to be released in the wild. The orangutan need to forage, build their own nest, and subsequently making them less dependent on human care. Nest building skills is a very crucial and important skill for orangutan to survive in the wild. Therefore, research on the nest structure of orangutan is a very important. We propose that our method would be used as a standard for future nest studies.

## Acknowledgement

We thank Universiti Sains Malaysia (USM) for providing the Research University grant. We also want to thank Sabah Biodiversity Centre (SABC) for approval of research permit. Thank you also to Sepilok Orangutan Rehabilitation Centre (SORC); Mr. Sailun, Mrs Sylvia and SORC staffs for assisting us during the sampling period. We acknowledge the Department of Aviation Malaysia (DCA) for the drone flight permission. A special thank you to Muhamad Armi Majid and Muhammad Shadzmir Kamalrulzaman from OFO Tech Snd. Bhd. for technical and drone piloting. Finally, our gratitude to Nagao Natural Environment Foundation, Orangutan Foundation International for the additional funding.

## References

1. Prasetyo D, Utami SS, Suprijatna J. Nest stuctures in Bornean Orangutan. Jurnal Biologi Indonesia. 2012;8(2).

2. Russon AE, Handayani DP, Kuncoro P, Ferisa A. Orangutan leaf-carrying for nest-building: Toward unraveling cultural processes. Animal cognition. 2007;10(2):189–202.

3. Russon AE. Orangutan rehabilitation and reintroduction. Orangutans: Geographic variation in behavioral ecology and conservation. 2009:327–50.

4. Utami Atmoko S, Arif Rifqi M. Buku panduan survei sarang orangutan: Forum Orangutan Indonesia & Fakultas Biologi Universitas Nasional; 2012.

5. Prasetyo D, Ancrenaz M, Morrogh-Bernard HC, Utami Atmoko S, Wich SA, van Schaik CP. Nest building in orangutans. Orangutans: Geographical Variation in Behavioral Ecology, Oxford University Press, Oxford. 2009:269–77.

6. Samson DR, Hunt KD. Chimpanzees preferentially select sleeping platform construction tree species with biomechanical properties that yield stable, firm, but compliant nests. PloS one. 2014;9(4):e95361.

7. van Casteren A, Sellers WI, Thorpe SK, Coward S, Crompton RH, Myatt JP, et al. Nest-building orangutans demonstrate engineering know-how to produce safe, comfortable beds. Proceedings of the National Academy of Sciences. 2012;109(18):6873–7.

8. Ancrenaz M, Gimenez O, Ambu L, Ancrenaz K, Andau P, Goossens B, et al. Aerial surveys give new estimates for orangutans in Sabah, Malaysia. PLoS Biology. 2004;3(1):e3.

9. Wich SA, Boyko RH. Which factors determine orangutan nests’ detection probability along transects? Tropical Conservation Science. 2011;4(1):53–63.

10. Mathewson P, Spehar S, Meijaard E, Sasmirul A, Marshall AJ. Evaluating orangutan census techniques using nest decay rates: Implications for population estimates. Ecological Applications. 2008;18(1):208–21.

11. Koh LP, Wich SA. Dawn of drone ecology: low-cost autonomous aerial vehicles for conservation. Tropical Conservation Science. 2012;5(2):121–32.

12. Ancrenaz M, Calaque R, Lackman-Ancrenaz I. Orangutan nesting behavior in disturbed forest of Sabah, Malaysia: Implications for nest census. International Journal of Primatology. 2004;25(5):983–1000. doi: 10.1023/B:IJOP.0000043347.84757.9a.

13. Rayadin Y, Saitoh T. Individual variation in nest size and nest site features of the Bornean orangutans (*Pongo pygmaeus*). American Journal of Primatology. 2009;71(5):393–9.

14. Houle A, Chapman CA, Vickery WL. Tree climbing strategies for primate ecological studies. International Journal of Primatology. 2004;25(1):237–60.

15. van Casteren A, Sellers W, Thorpe S, Coward S, Crompton R, Ennos A. Why don’t branches snap? The mechanics of bending failure in three temperate angiosperm trees. Trees. 2011;26(3):789–97.

16. Kindervater KH. The emergence of lethal surveillance: Watching and killing in the history of drone technology. Security Dialogue. 2016;47(3):223–38.

17. Floreano D, Wood RJ. Science, technology and the future of small autonomous drones. Nature. 2015;521(7553):460–6.

18. Heatherly MC. Drones: The American controversy. Journal of Strategic Security. 2014;7(4):25.

19. Paneque-Gálvez J, McCall MK, Napoletano BM, Wich SA, Koh LP. Small drones for community-based forest monitoring: An assessment of their feasibility and potential in tropical areas. Forests. 2014;5(6):1481–507.

20. Flynn KF, Chapra SC. Remote sensing of submerged aquatic vegetation in a shallow non-turbid river using an unmanned aerial vehicle. Remote Sensing. 2014;6(12):12815–36.

21. Sugardjito J. Selecting nest-sites of Sumatran organ-utans, *Pongo pygmaeus abelii* in the Gunung Leuser National Park, Indonesia. Primates. 1983;24(4):467–74.

22. Koops K, McGrew WC, de Vries H, Matsuzawa T. Nest-building by chimpanzees (*Pan troglodytes verus*) at Seringbara, Nimba Mountains: Antipredation, thermoregulation, and antivector hypotheses. International Journal of Primatology. 2012;33(2):356–80.

23. Mulyoutami E, Rismawan R, Joshi L. Knowledge and use of local plants from ‘simpukng’or forest gardens among the Dayak community in East Kalimantan. IUFRO World Series Vol 21. 2008:121.

24. IUCN. The IUCN Red List of Threatened Species 2018 [cited 2017 13th December]. Available from: http://www.iucnredlist.org

25. Zahorka H. The Shamanic belian sentiu rituals of the Benuaq Ohookng, with special attention to the ritual use of plants. Borneo Research Bulletin. 2007;38:127–47.

26. Pereira JT, Sugau JB. A Guide to the Trees in Heritage Amenity Forest Reserve, Sandakan: Sabah Forestry Department; 2010.

27. Cheyne SM, Rowland D, Höing A, Husson SJ. How orang-utans choose where to sleep: Comparison of nest site variables. Asian Primates Journal. 2013;3:13–7.

28. Delgado RA, van Schaik CP. The behavioral ecology and conservation of the orangutan (Pongo pygmaeus): A tale of two islands. Evolutionary Anthropology: Issues, News, and Reviews. 2000;9(5):201–18.

29. Rijksen HD. A fieldstudy on Sumatran Orang Utans (Pongo pygmaeus abelii Lesson 1827): Ecology, behaviour and conservation: Veenman; 1978.

